# Isolation and characterization of a novel phage of *Vibrio parahaemolyticus* with biocontrol potential

**DOI:** 10.1101/2022.11.25.517947

**Authors:** Yubing Chen, Wenqing Li, Keming Shi, Zheng Fang, Yunlan Yang, Rui Zhang

## Abstract

*Vibrio parahaemolyticus* is a major foodborne pathogen that contaminates aquatic products and causes great economic losses to aquaculture. Because of the emergence of multidrug-resistant *V. parahaemolyticus* strains, bacteriophages are considered promising agents for their biocontrol as an alternative or supplement to antibiotics. Here, a lytic vibriophage, vB_VpaM_R16F (R16F), was isolated from sewage from a seafood market by infecting *V. parahaemolyticus* 1.1997^T^. R16F was found to infect *V. parahaemolyticus*, but not nine other *Vibrio* spp. The phage belongs to the myovirus morphotype and lysed host cells with a short latent period (<10 min) and a small burst size (13 plaque-forming units). R16F has a linear double-stranded DNA with genome size 139,011 bp and a G+C content of 35.21%. Phylogenetic and intergenomic nucleotide sequence similarity analysis revealed that R16F is distinct from currently known vibriophages and belongs to a novel genus. Several genes (e.g., encoding ultraviolet damage endonuclease and endolysin) that may enhance environmental competitiveness were found in the genome of R16F, while no antibiotic resistance- or virulence factor-related gene was detected. In consideration of its biological and genetic properties, R16F is suggested to be a candidate biocontrol agent for use against *V. parahaemolyticus*.

## 1. Introduction

Vibriosis is one of the most prevalent bacterial diseases that causes mortality of shrimp, fish and shellfish. It results from contamination with *Vibrio* pathogens, such as *V. parahaemolyticus, V. vulnificus, V. alginolyticus* and *V. harveyi* [1,2]. With the rapid development of aquaculture and the rise in consumption of aquatic products, vibriosis in global aquaculture and *Vibrio*-related food poisoning cases are increasing [1,3,4]. *V. parahaemolyticus* is ubiquitous in seawater and seafood-associated environments [5–7]. According to a comprehensive review and meta-analysis of studies between 2003 and 2015 on the occurrence and prevalence of *V. parahaemolyticus* in seafood, it could be isolated from 47.5% of all seafood samples, and showed high overall prevalence rates in oysters (63.4%), clams (52.9%), fish (51.0%), shrimps (48.3%), and mussels, scallop and periwinkle (28.0%) [8]. Eating any raw *V. parahaemolyticus*-contaminated seafood can lead to gastroenteritis because the pathogen contains various virulence factors, including adhesin, heat-resistant direct hemolysin (TDH), TDH-associated hemolysin, and the type III secretory system [9]. The reported incidence of *V. parahaemolyticus* in fishery products in China was approximately 15% during the period 2008 to 2017 [10]. The administration of antibiotics to quickly and effectively control pathogenic *Vibrio* strains became common in aquaculture. However, prolonged use of antibiotics has resulted in an increase in the number of multidrug-resistant *Vibrio* strains, presenting a substantial threat to the control of vibriosis [11–13]. Calls have been made to decrease antibiotic use, and alternative measures to control bacterial pathogens are urgently required.

Phages infect and lyse host bacterial cells and plunder host cell resources for their own reproduction [14–17]. Phages have been known as a potential antibacterial agent for over a century; they were used against human bacterial infection in the 1920s [18]. Previous studies have reported on global efforts to prevent and control *Vibrio* in aquaculture by using phages. Numerous *in vitro* and *in vivo* studies have shown that phage therapy was effective in controlling vibriosis caused by *Vibrio* (e.g., *V. parahaemolyticus* and *V. harveyi*) in shrimp, sea cucumber, abalone, oysters, and other species [19–23]. By way of illustration, a 78.1% reduction in bacterial counts was observed within 1 h of phage application during an efficacy study of phage against *V. parahaemolyticus* in shrimp [23]. *V. parahaemolyticus*-infected shrimp larvae regained activity with no significant reduction in survival when they were treated with phage [20]. The application of phages in a variety of aquaculture situations highlights the potential of phage therapy to decrease bacterial infections and diminish the significant economic losses to aquaculture and the harm to public health caused by contamination with *Vibrio*.

The selection of appropriate phages is, however, a prerequisite, and important tests need to be performed before phage therapy can be applied in the field. In this study, a novel bacteriophage that infects *V. parahaemolyticus* 1.1997^T^ was isolated from sewage from the Chigang seafood market in Guangdong, China. The phage was characterized and evaluated for possible use in phage therapy.

## 2. Results and Discussion

### 2.1 Biological properties of phage R16F

Vibriophage vB_VpaM_R16F (hereafter R16F) was isolated from a sewage sample from Chigang seafood market, Guangzhou, China, by infecting *V. parahaemolyticus* 1.1997^T^. R16F formed clear plaques around 2–3 mm in diameter after infection for 12 h (Figure 1A). With extension of the incubation time, the phage plaques became larger (up to 7–8 mm in diameter) with a turbid halo on the host bacterial lawn (Figure 1A). Halo formation has previously been described as an indicator of phage-associated exopolysaccharide depolymerization [24,25]. The depolymerases encoded by phages can specifically cleave polysaccharides (e.g. capsular polysaccharides, exopolysaccharides or lipopolysaccharides) of the host bacteria and therefore provide a significant advantage for phages to adsorb onto their hosts [26,27]. Such halo zones have been observed for various phages, including phages that infect *Pseudomonas putida, Klebsiella pneumoniae*, and *V. alginolyticus* [28–30]. TEM revealed that R16F has a narrow neck or collar region, an icosahedral capsid (head diameter 83 ± 7 nm), and a complete tail with contracted sheath (49 ± 2–100 ± 3 nm). It belongs to the myovirus morphotype (Figure 1B).

**Figure 1.**
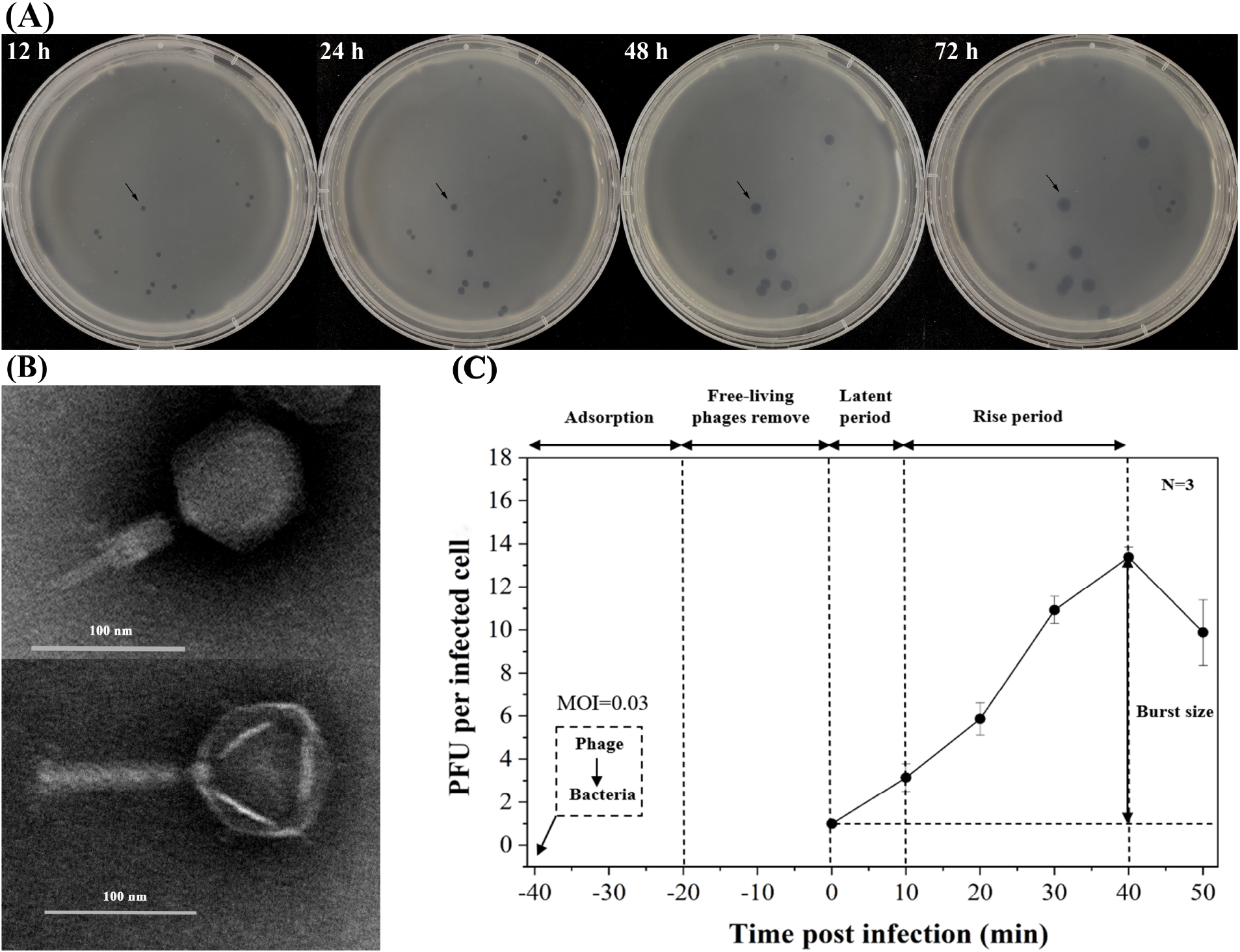
Morphology and infection dynamics of vibriophage vB_VpaM_R16F (R16F). (A) The plaque morphology of vibriophage R16F after incubation with *V. parahaemolyticus* 1.1997^T^ for 12, 24, 48 and 72 h. The phage plaques are surrounded by expanding opaque haloes as the incubation time is extended. (B) Transmission electron micrographs of vibriophage R16F with contracted tail (upper panel) and non-contracted tail (lower panel). (C) One-step growth curve of vibriophage R16F. Phages were incubated with an early log-phase *V. parahaemolyticus* 1.1997^T^ at a multiplicity of infection of 0.03. PFU, plaque-forming units.

As Figure 1C shows, R16F exhibited a short latent period, <10 min, and then reached the plateau within 40 min of infection. The burst size for R16F, defined as the total number of progeny virions liberated by one infected host cell at the completion of a growth cycle, estimated from the one-step growth curve, was 13 plaque-forming units (PFU) cell^−1^ (Figure 1C). Short latent periods (<30 min) and small burst sizes (ranging from 10 to 100 PFU cell^−1^) have been observed for other phages that infect *V. parahaemolyticus* [7,31–39]. A trade-off between burst size and latent period has been reported in lytic phage generation because phages are released from infected cells at the cost of destruction of the machinery necessary to produce more phage progeny [40]. Thus, phages with short latent periods at the cost of their burst size may be viewed as specialists for propagation when bacteria are prevalent.

R16F exhibited a narrow lytic spectrum and only infected the original host *V. parahaemolyticus* 1.1997^T^ among the species tested (Table S1), which included strains of 10 *Vibrio* spp. R16F was insensitive to chloroform, suggesting that there was no lipid in the viral capsid. This result was consistent with the previous suggestion that lipids are rare among phages, occurring in <4% of isolates [41].

### 2.2 Overview of genomic features and taxonomy of R16F

Whole genome sequencing revealed that the R16F genome is a 139,011-bp linear double-stranded DNA with a G+C content of 35.21%, which is lower than its host (45.3%). According to the results from PhageTerm, a fixed position was recognized on the DNA of R16F and 353-bp direct terminal repeats (DTR) were observed, suggesting that the DNA termini of R16F belong to the DTR class. A total of 260 ORFs were identified in the genome of R16F, of which only 57 (21.92%) have known functional domains, while the remaining 203 (70.77%) were annotated as hypothetical proteins (Figure 2; Table S2). The functional ORFs were identified to encode proteins that can be arranged in functional categories including phage structural formation, genome packaging, nucleotide metabolism, integration, lysis, and other functions. Among these functional ORFs, about 12.69% (33/260) are conserved. The genome of R16F was predicted to encode two tRNAs (*tRNA*^*Arg*^ and *tRNA*^*Met*^) carried anticodons TCT (serine) and CAT (histidine). No antibiotic resistance- or virulence factor-related gene was found in the genome of R16F.

**Figure 2.**
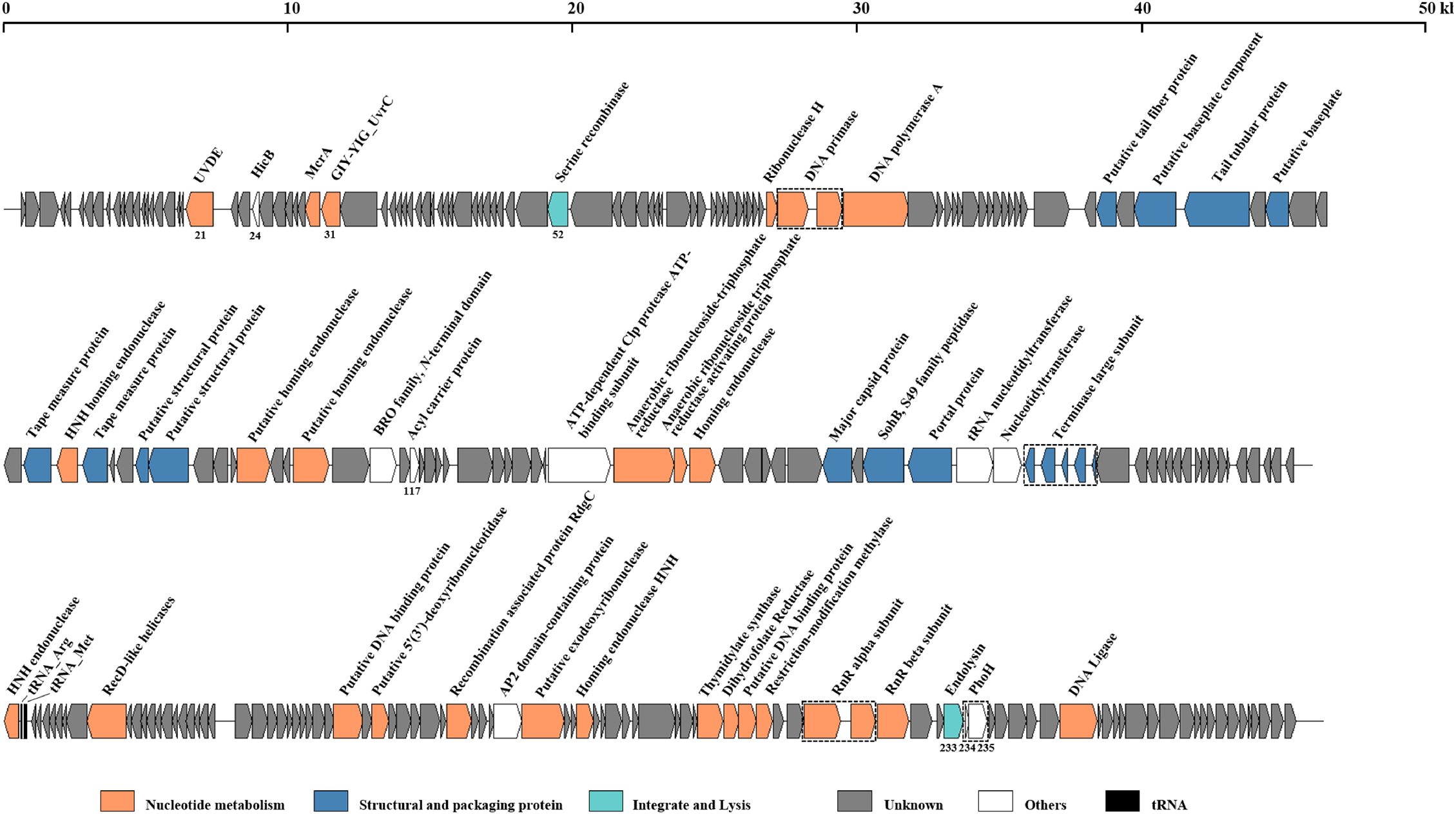
Genomic organization of vibriophage R16F. Putative open reading frames are represented by arrows and were classified into different functional categories indicated by the colors shown at the bottom of the figure. UVDE, putative UV DNA damage repair endonuclease; HicB, type II toxin–antitoxin system HicB family antitoxin; McrA, 5-methylcytosine-specific restriction endonuclease; GIY-YIG UvrC, catalytic GIY-YIG domain of nucleotide excision repair endonucleases UvrC, Cho, and similar proteins; PhoH, phosphate starvation-inducible protein.

On the basis of NCBI BLASTN analysis, phages qdvp001, VPMCC14, and PWH3a-P1 share the highest sequence identities (95.15%, 84.98%, and 88.33%, respectively) with R16F, but they share only 33%, 15%, and 12% query coverage. To investigate the evolutionary relationships and taxonomic status of R16F, the vConTACT2 database was used to detect other homologous phages. A total of 37 phages with a similarity score of >1 were identified, including six vibriophages with high scores of >40 (Table S3). All the phages detected by vConTACT2 and other phage sequences obtained from NCBI BLASTN were used to construct a phylogenetic tree based on pairwise comparisons of the amino acid sequences of R16F. According to the phylogenetic analysis, R16F was most closely related to vibriophages qdvp001, VPMCC14 and PWH3a-P1 (Figure 3A). To better estimate the similarity between phage genomes, all phages from the tree were imported into the VIRIDIC software and demarcated based on the default similarity thresholds of 95% for species and 70% for genus [42,43]. The VIRIDIC results showed that the highest value of intergenomic similarity was only 34.9%, observed between R16F and qdvp001 (Figure 3B). Therefore, R16F is suggested to belong to a novel genus. Overall, R16F is considered a novel phage with a genome that differs from all previously described phage sequences.

**Figure 3.**
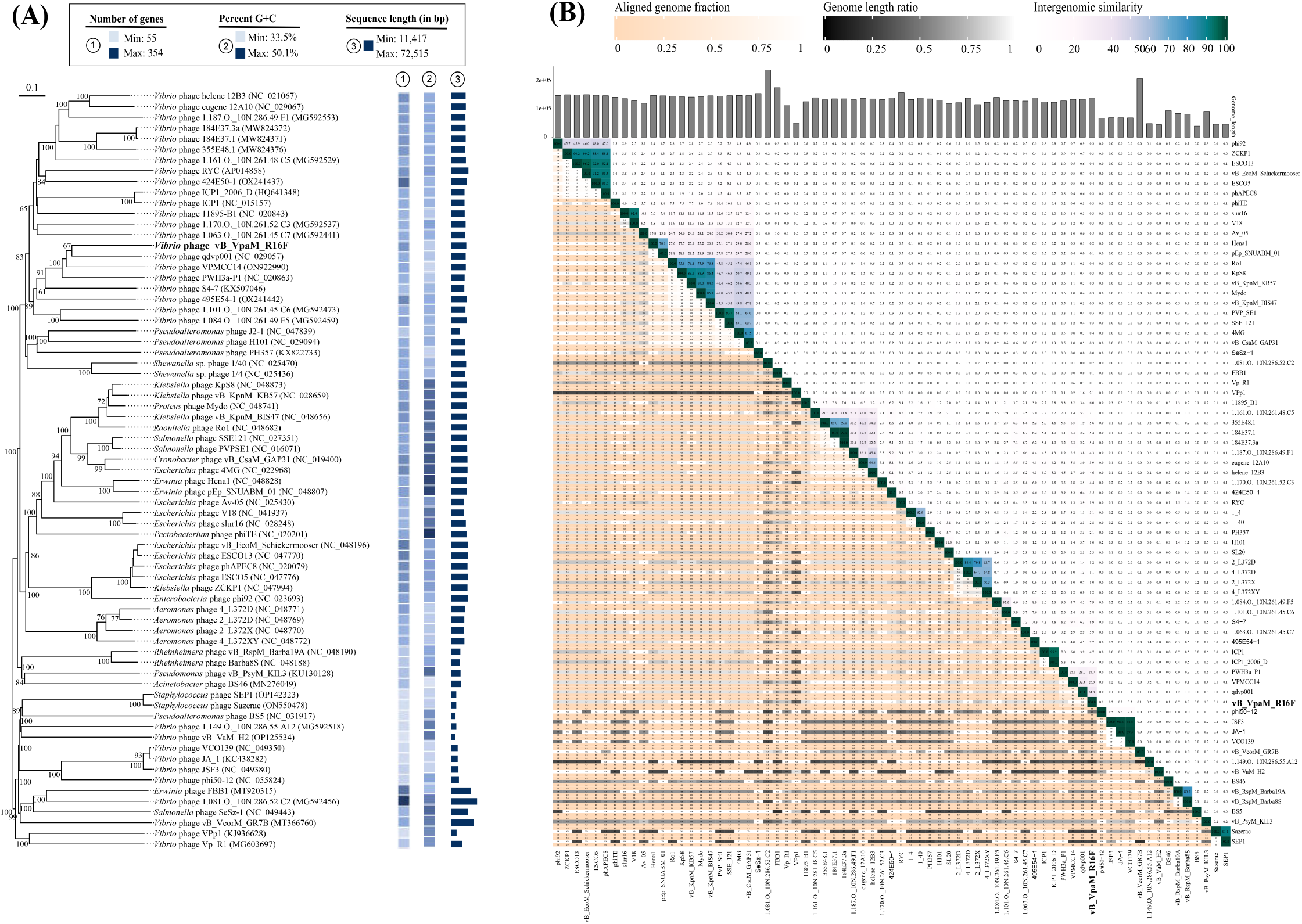
Taxonomy of phage R16F. (A) Phylogenetic tree, including R16F and other closely related phages, constructed using the Virus Classification and Tree Building Online Resource (VICTOR) web service. Pairwise comparisons of the amino acid sequences were conducted using the Genome-BLAST Distance Phylogeny (GBDP) method with settings recommended for prokaryotic viruses. The numbers above branches are GBDP pseudo-bootstrap support values from 100 replications. (B) Virus Inter-genomic Distance Calculator (VIRIDIC)-generated heatmap incorporating intergenomic similarity values (right half) and alignment indicators (left half and top annotation).

### 2.3 DNA repair genes in the genome of phage R16F

Phage R16F harbors a gene (ORF 21) encoding an ultraviolet damage endonuclease (UVDE), which is involved in DNA repair. UV light may cause damage or mortality to bacteria and phages [44]. In bacterial cells, UVDE has been defined as a repair protein that is capable of identifying and removing various DNA lesions including UV photoproducts as well as non-UV-induced DNA damage such as abasic sites, strand breaks, and gaps [45]. In addition to being found in the genomes of bacteria and fungi, UVDE is present in the genomes of some phages [46,47]. UVDE of bacteriophage T4 was involved in the dark repair of UV light damage in T4-infected cells [48]. Furthermore, a conserved catalytic GIY-YIG domain of nucleotide excision repair endonucleases UvrC, Cho, and similar proteins, is encoded in the R16F genome (ORF 31). UvrA, UvrB, and UvrC mediate nucleotide excision repair following various types of structurally unrelated DNA damage and remove the complete oligonucleotide containing the damage [49–53].

### 2.4 Type II toxin-antitoxin (TA) system HicB family antitoxin gene

A gene (ORF 24) encoding a type II TA system HicB family antitoxin (HicB) was found in the genome of R16F. TA systems are composed of a two-gene cassette encoding a toxic protein and an antitoxin protein, the latter counteracting the toxicity of the former [54]. Type II TA systems appear to be the most abundant and diverse in bacterial and archaeal genomes, largely being found in mobile gene modules such as plasmids and phages, but also in bacterial chromosomes, and they are highly prone to horizontal gene transfer [54–56]. Type II TA systems in bacteria are thought to be involved in diverse biological processes, including plasmid maintenance, phage inhibition, persistence, stress response, and biofilm formation [54,57]. The *hicAB* system, a type II TA system, encoding toxin HicA and antitoxin HicB proteins, has been predicted to be associated with several functions, including RNA-binding, persister cell formation and involvement in extracytoplasmic stress responses [55,58,59]. On the basis of BLASTP analysis, almost all homologues of the HicB in R16F were from bacterial genomes, implying that the HicB in R16F most likely derived from bacteria via horizontal gene transfer. However, no gene encoding a HicA toxin protein was identified in R16F. Hitherto, intact TA systems have been found in prophages, such as prophages induced from *Streptococcus suis, Mannheimia haemolytica* and *Acinetobacter*. However, this is the first report of a component of a TA system in a lytic phage [60–62]. A recent review suggested that phages have evolved defenses against TA systems by incorporating active TA systems [63]. The roles of TA systems in phage genomes, especially those of lytic phages, remain unclear, but they are speculated to perform functions in phage–bacteria co-evolution.

### 2.5 Acyl carrier protein (ACP)

ACP (ORF 117 in phage R16F) serves as a ubiquitous and highly conserved transporter of acyl intermediates during the synthesis of fatty acids [64]. ACPs in bacteria are acyl donors for synthesis of various products, including endotoxin and acylated homoserine lactones, which are involved in quorum sensing. The unique nature of these growth and pathogenesis processes makes ACP-dependent enzymes a target for antibacterial agents [64]. In addition to being widespread in bacteria, homologs of ACP have been identified in several phages, such as *V. helene* phage 12B3, *Erythrobacter* phage vB_EliS-R6L, and some prophages [65,66]. ACP encoded by phages was previously identified as an auxiliary metabolic gene and potentially involved in the biosynthesis of secondary metabolites in host cells [66].

### 2.6 Phosphate starvation-inducible protein (phoH)

The genome of phage R16F also harbors a *phoH* gene (ORFs 234 and 235). In *Escherichia coli*, gene transcription of the phosphate regulator is activated in response to phosphate restriction, and its products participate in the transportation and use of various forms of phosphate [67–69]. The *phoH* gene has been detected in marine phages, including vibriophages, cyanophages, and roseophages [70-74]. Although many studies have hypothesized that phage-encoded *phoH* might play an important role in assisting host phosphorous metabolism, no concrete evidence has been reported to support this conjecture.

### 2.7 Phage therapy potential inferred from biological and genomic features of R16F

The biological and genomic characteristics are critical in evaluating phage fitness and identifying candidates for use in phage therapy. Several general properties should be considered, including bactericidal ability, efficiency, and whether the phage carries endotoxins [75]. Phages have often evolved to quickly suppress bacterial proliferation when the bacterial density is sufficiently high [40]. This phenomenon was observed here for R16F (latent period <10 min, with small burst size). Therefore, R16F may be effective against *Vibrio* outbreaks. The narrow lytic spectrum (host range) of R16F would enable use of this phage to specifically target *V. parahaemolyticus*. Moreover, previous study suggested that various type of phage with narrow host specificity can be mixed in cocktails to control a range of pathogenic strains [32].

R16F was found to encode an endolysin (ORF 233) that would be able to hydrolyze the bacterial peptidoglycan layer of the host cell wall and cause lysis of the host bacterium and release newly assembled phage particles [76]. Phage-encoded endolysin is also suggested to facilitate cell adhesion of phages by degrading the capsular polysaccharides, and phages encoding endolysin are believed to have good performance against bacterial biofilms [26,77]. The presence of the endolysin in the R16F genome is consistent with the halo zone observed around plaques (Figure 1A). The R16F-encoded endolysin showed 95.76% amino acid sequence identity with the endolysin encoded by phage qdvp001 (lysqdvp001). Lysqdvp001 and its homologues are highly divergent enzymes with superior lytic activity and a broader spectrum than their parent phages [35,78]. Therefore, both vibriophage R16F and R16F-derived endolysin are promising for application in biocontrol.

No antibiotic resistance- or virulence factor-related gene was predicted in the genome of R16F using VFDB and CARD, indicating that R16F infection is unlikely to result in enhancement of *Vibrio* virulence or contamination of aquaculture environments with antibiotic resistance genes. Although R16F encodes a serine recombinase (ORF 52), no attachment sites on the phage or bacterial genomes (*attP–attB*) were observed in the 1000-nt sequences adjacent to either end of the serine recombinase-encoding gene. We postulate that R16F is likely to follow a lytic lifestyle. Therefore, R16F is considered safe at the genetic level as a biocontrol agent.

## 3. Materials and Methods

### 3.1 Phage isolation and purification

*V. parahaemolyticus* 1.1997^T^ brought from Guangdong marine pathogenic *Vibrio* company in China was used as the host bacterium for phage isolation. It was incubated in rich organic (RO) medium (1 M peptone, 1 M yeast extract, and 1 M sodium acetate in artificial seawater, pH 7.5) at 30°C with a shaking speed of 160 rpm min^-1^ [70]. The sewage samples used for phage isolation were collected from the seafood markets in Guangzhou, China (23.10°N, 113.33°E), and filtered through a 0.22-μm membrane (Millipore, Massachusetts, USA) to remove bacteria and large particles. The sewage samples were added into an exponentially growing host bacterial culture to allow plaque formation using the double-layer agar method [79]. A clear individual plaque was collected and suspended in storage medium (SM; 8 mM MgSO_4_, 50 mM Tris-HCl, and 100 mM NaCl, pH 7.5). After purifying at least five times, the well-separated plaque was collected and stored in SM at 4°C.

### 3.2 Phage amplification and enrichment

To obtain high-titer phage suspension, a purified phage plaque was inoculated into bacterial culture and amplified overnight, followed by centrifugation at 12,000 × *g* for 10 min. The supernatant was filtered through a 0.22-μm membrane to remove cell fragments. The filtrate was precipitated with polyethylene glycol 8000 (10% w/v) overnight. Then, phage pellets were obtained through centrifugation (10,000 × *g*, 60 min, 4°C) and resuspended in SM. The phage particles were subjected to cesium chloride solutions (ρ = 1.3, 1.5, 1.7 g mL^−1^) for purification and centrifuged at 200,000 × *g* at 4°C for 24 h using an Optima L-100 XP ultracentrifuge (Beckman Coulter, CA, USA). The purified phage particles were dialyzed through 30-kDa superfilters (Millipore, Bedford, MA, USA).

### 3.3 Transmission electron microscopy (TEM)

Phage morphology was characterized by TEM. Briefly, approximately 20 μL of phage suspension was added onto the surface of a copper grid to adsorb in darkness for 30 min. Then, the phage sample was negatively stained with phosphotungstic acid (1%, pH 7.0) for 20 min and dried for 30 min. The phage sample was examined using a JEM-2100 transmission electron microscope (JEOL, Tokyo, Japan).

### 3.4 Determination of the host range

A total of 22 *Vibrio* strains, including seven strains of *V. parahaemolyticus* and 15 strains of other species (*V. alginolyticus, V. campbellii, V. caribbeanicus, V. cholerae, V. fortis, V. harveyi, V. inhibens, V. owensii*, and *V. variabilis*), were used to determine the lytic host range of the phage isolated in this study (Table S1). The double-layer agar method was used for host-range tests with two replicates. The presence of plaques on a bacterial lawn was checked to determine phage infection of the host bacterium.

### 3.5 Lipid test

To investigate whether the phage capsids contain lipid, phages were mixed with 0%, 0.2%, 2% or 20% chloroform and incubated in darkness at room temperature for 30 min. Then, the mixtures were centrifuged at 12,000 × *g* for 5 min and the phages were obtained from the upper suspension. The chloroform sensitivity of the phages was determined by the presence or absence of plaqueson double-layer agar. The lipid test was carried out twice.

### 3.6 One-step growth curve

A one-step growth curve was determined to study the infectivity and replication ability of the phage. Briefly, phages were added to an early log-phase host culture (*V. parahaemolyticus* 1.1997^T^) at a multiplicity of infection of 0.03 and incubated for 20 min at room temperature in the dark. Then, phages that were not adsorbed were removed by centrifugation (8,000 × *g*, 4°C, 5 min), and the cell pellets were washed and resuspended in 100 mL of RO medium. The phage suspension was incubated at 28°C with a shaking of 160 rpm min^-1^. Subsamples were obtained at 10-min intervals and assayed by the double-layer agar method. The burst size was calculated as the ratio between the number of phages before and after the burst [80].

### 3.7 DNA extraction and phage genome sequencing

Phage DNA was obtained by using the phenol–chloroform extraction method. Briefly, approximately 1 mL of high-titer phage suspension was treated with proteinase K, sodium dodecyl sulfate (10% w/v), and ethylenediaminetetraacetic acid(EDTA; pH 8.0) and incubated at 55°C for 3 h. The digested sample was purified with phenol/chloroform/isoamyl alcohol (25:24:1 v:v:v) and chloroform/isoamyl alcohol (24:1 v:v) to remove any debris. Next, the DNA pellet was washed with pre-cooled 70% ethanol, air-dried at room temperature, dissolved in Tris-EDTA buffer (10 mM Tris-HCl, 1 mM EDTA, pH 8.0), and stored at 4°C. The phage genome was sequenced and assembled using the Illumina HiSeq 4000 platform with a 150-bp paired-end DNA library and Velvet software (v.1.2.03) [81].

### 3.8 Genome annotation

The phage termini and packaging mechanism were identified by the online service PhageTerm on a Galaxy-based server (https://galaxy.pasteur.fr) [82]. The GeneMarkS online server (http://topaz.gatech.edu/GeneMark/genemarks.cgi) was used to identify putative open reading frames (ORFs) [83]. The functions of ORFs were annotated by BLASTP online against the non-redundant (nr) database of the National Center for Biotechnology Information (NCBI), with a cut-off *e*-value of <10^−5^, and the results were checked manually. Putative transfer RNA (tRNA) genes were detected with tRNAscan-SE (http://lowelab.ucsc.edu/tRNAscan-SE/) [84]. Virulence factors and antibiotic resistance-encoding genes were searched using the Virulence Factor Database (VFDB, http://www.mgc.ac.cn/VFs/main.htm) and the Comprehensive Antibiotic Resistance Database (CARD, https://card.mcmaster.ca/analyze/rgi) webservers, respectively [85,86]. The complete genome data was deposited in the GenBank database with accession number OP793884.

### 3.9 Phylogenetic and taxonomic analysis

Viral CONTigs Automatic Clustering and Taxonomy v.2.0 (vConTACT2) was used to compare the isolated phage against the Prokaryotic Viral RefSeq 207 database using whole genome gene-sharing profiles, and related phages were identified by genome pairs with a similarity score >1 [87]. The whole phage sequence was also submitted to NCBI BLASTN to search forsimilar sequences. To analyze the evolutionary relationships of the phage, complete amino acid profiles of phages were submitted to the Virus Classification and Tree Building Online Resource (VICTOR, https://ggdc.dsmz.de/victor.php) for phylogenetic tree construction [88,89]. All pairwise comparisons of the amino acid sequences were conducted using the Genome-BLAST Distance Phylogeny (GBDP) method with the settings recommended for prokaryotic viruses [88,90]. Branch support was inferred from 100 pseudo-bootstrap replicates each, and the tree was rooted at the midpoint and visualized with ggtree [91,92]. In addition, intergenomic nucleotide sequence similarity and aligned genome fractions within the imported phages were plotted with the Virus Inter-genomic Distance Calculator (VIRIDIC) using default parameters [42].

## 4. Conclusions

This study isolated a lytic phage, vB_VpaM_R16F (R16F), active against *Vibrio parahaemolyticus* 1.1997^T^ and investigated its biological properties. R16F is a myovirus with an icosahedral head and a contracted tail. R16F exhibits a narrow host spectrum, small burst size, and short latent period. A large portion of the genes of RF16 are annotated as hypothetical proteins. However, serval functional genes that may improve the environmental competitiveness of the phage were found in the genome of R16F. Antibiotic resistance or virulence factor-related genes were not detected. R16F is a newly described vibriophage, belonging to a novel genus. Its biological and genetic properties suggest that R16F has the potential to be used as a biocontrol agent for vibriosis induced by *V. parahaemolyticus*.

## Author Contributions

Conceptualization, R.Z., Y.Y. and Y.C.; methodology, R.Z., Y.Y., Y.C. and Z.F., W.L., K.S.; software, Y.Y. and W.L.; validation, R.Z., Y.Y., Y.C. and Z.F., W.L., K.S.; formal analysis, Y.C., Y.Y.; writing—original draft preparation, Y.C. and Y.Y.; writing—review and editing, R.Z., Y.Y. and Y.C.; supervision, R.Z. and Y.Y.; funding acquisition, R.Z. and Y.Y. All authors have read and agreed to the published version of the manuscript.

## Institutional Review Board Statement

Not applicable.

## Data Availability Statement

Not applicable.

## Acknowledgments

We are grateful to Dr. R.M. and Dr. L.W. for helpful advice on the manuscript.

## Supplementary materials

**Table S1.**
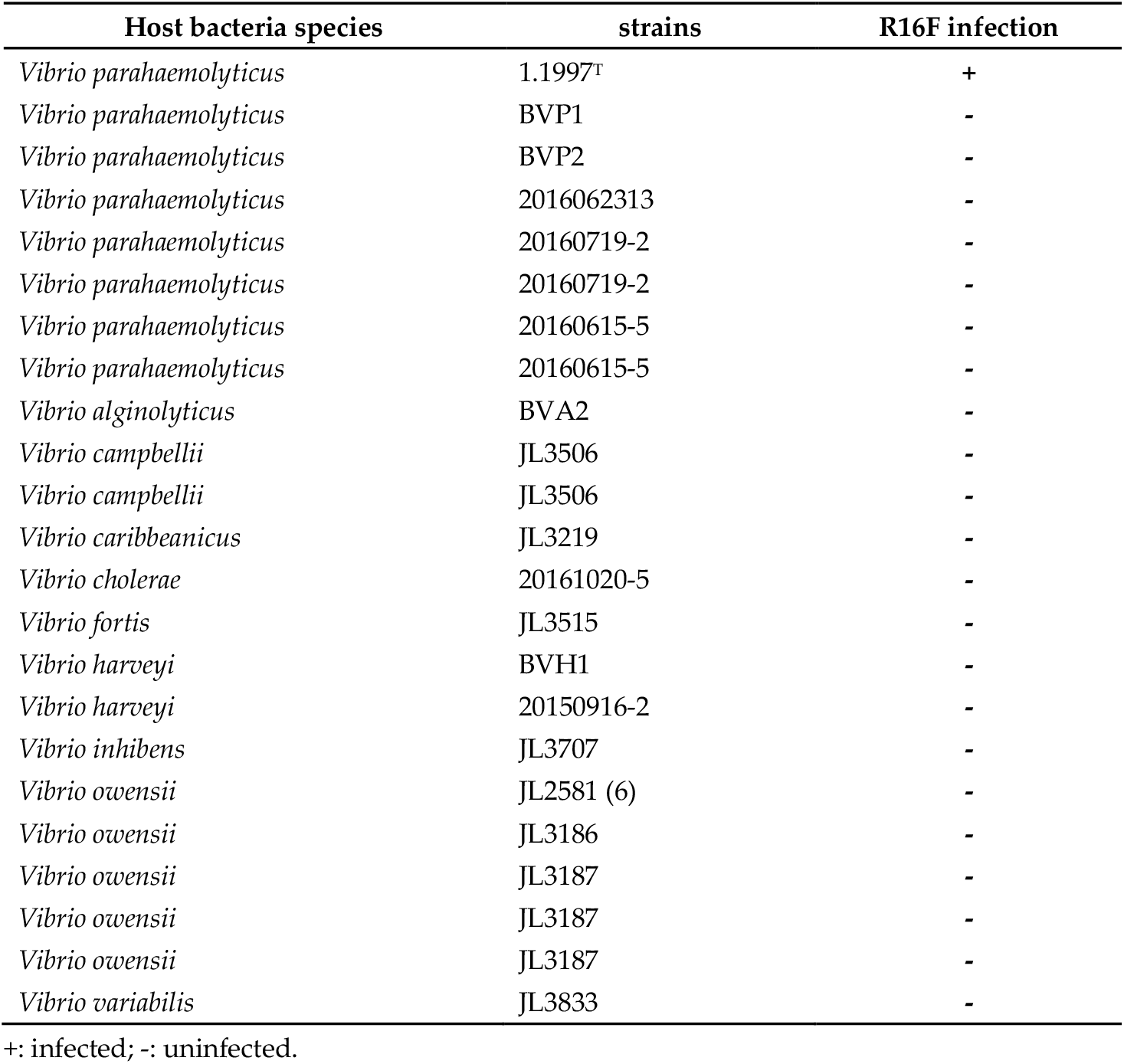
Host range of phage R16F.

| Host bacteria species | strains | R16F infection |
| --- | --- | --- |
| *Vibrio parahaemolyticus* | 1.1997<sup>T</sup> | + |
| *Vibrio parahaemolyticus* | BVP1 | - |
| *Vibrio parahaemolyticus* | BVP2 | - |
| *Vibrio parahaemolyticus* | 2016062313 | - |
| *Vibrio parahaemolyticus* | 20160719-2 | - |
| *Vibrio parahaemolyticus* | 20160719-2 | - |
| *Vibrio parahaemolyticus* | 20160615-5 | - |
| *Vibrio parahaemolyticus* | 20160615-5 | - |
| *Vibrio alginolyticus* | BVA2 | - |
| *Vibrio campbellii* | JL3506 | - |
| *Vibrio campbellii* | JL3506 | - |
| *Vibrio caribbeanicus* | JL3219 | - |
| *Vibrio cholerae* | 20161020-5 | - |
| *Vibrio fortis* | JL3515 | - |
| *Vibrio harveyi* | BVH1 | - |
| *Vibrio harveyi* | 20150916-2 | - |
| *Vibrio inhibens* | JL3707 | - |
| *Vibrio owensii* | JL2581 (6) | - |
| *Vibrio owensii* | JL3186 | - |
| *Vibrio owensii* | JL3187 | - |
| *Vibrio owensii* | JL3187 | - |
| *Vibrio owensii* | JL3187 | - |
| *Vibrio variabilis* | JL3833 | - |

+: infected; -: uninfected.

**Table S2.**
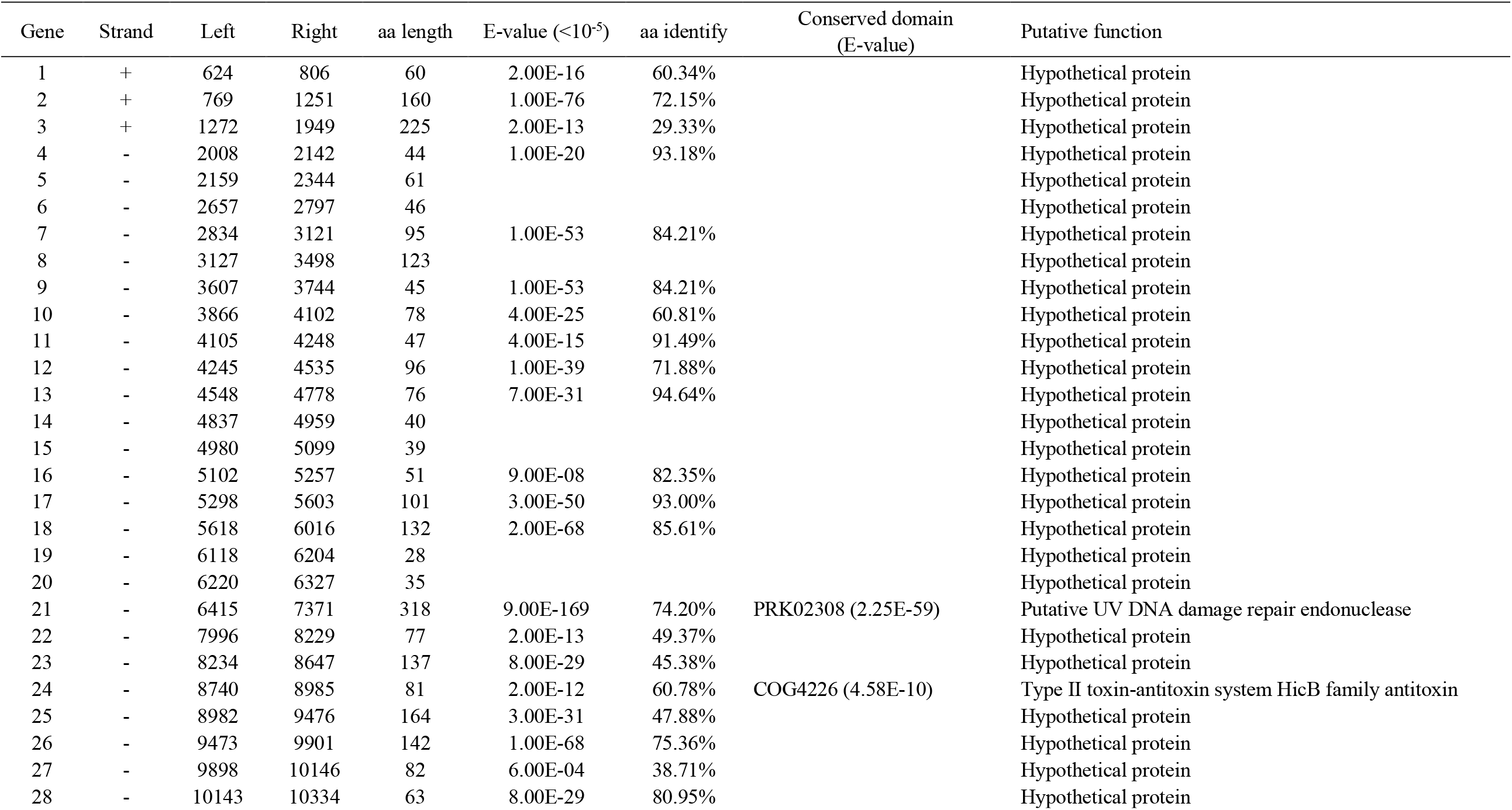
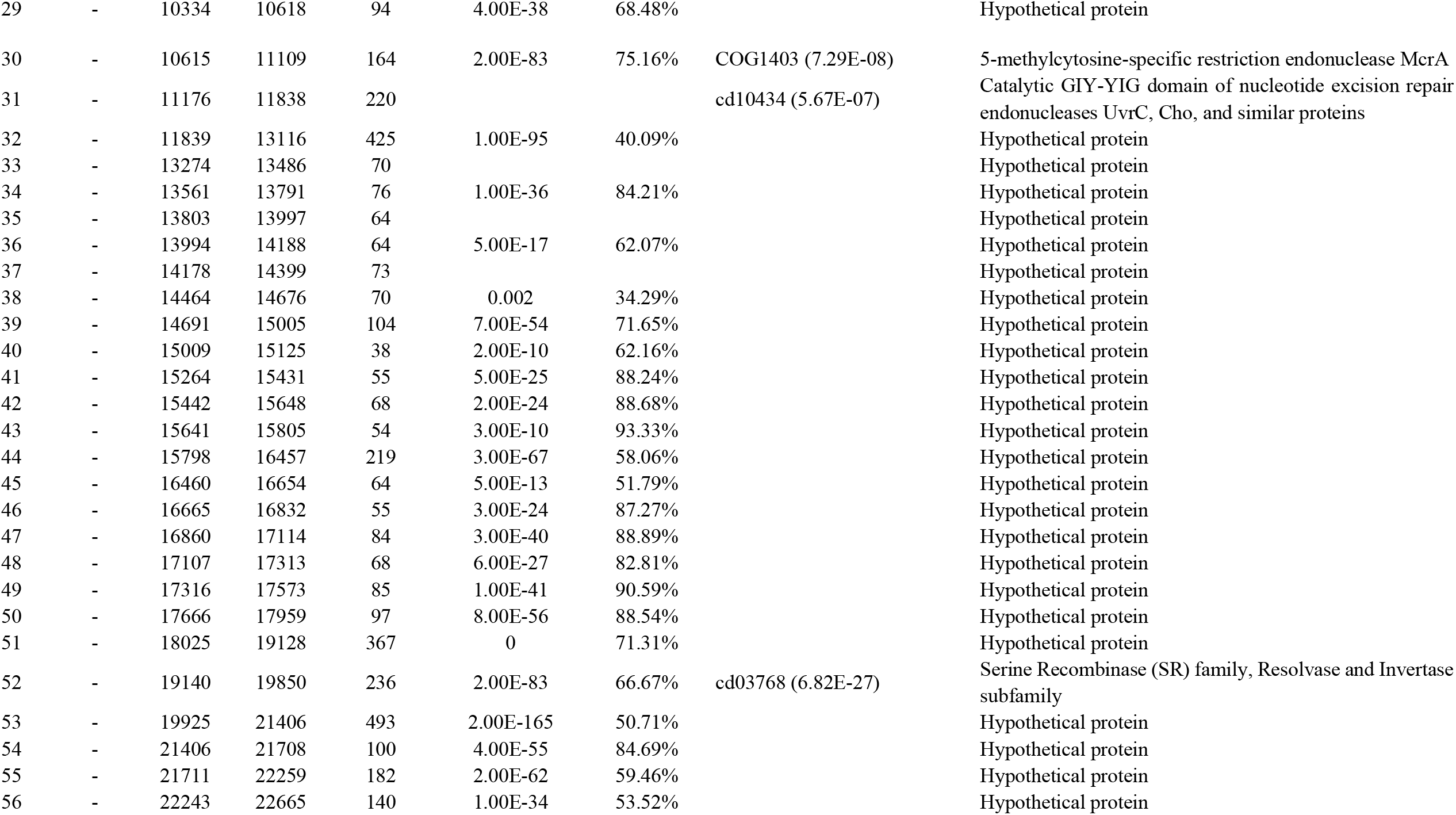
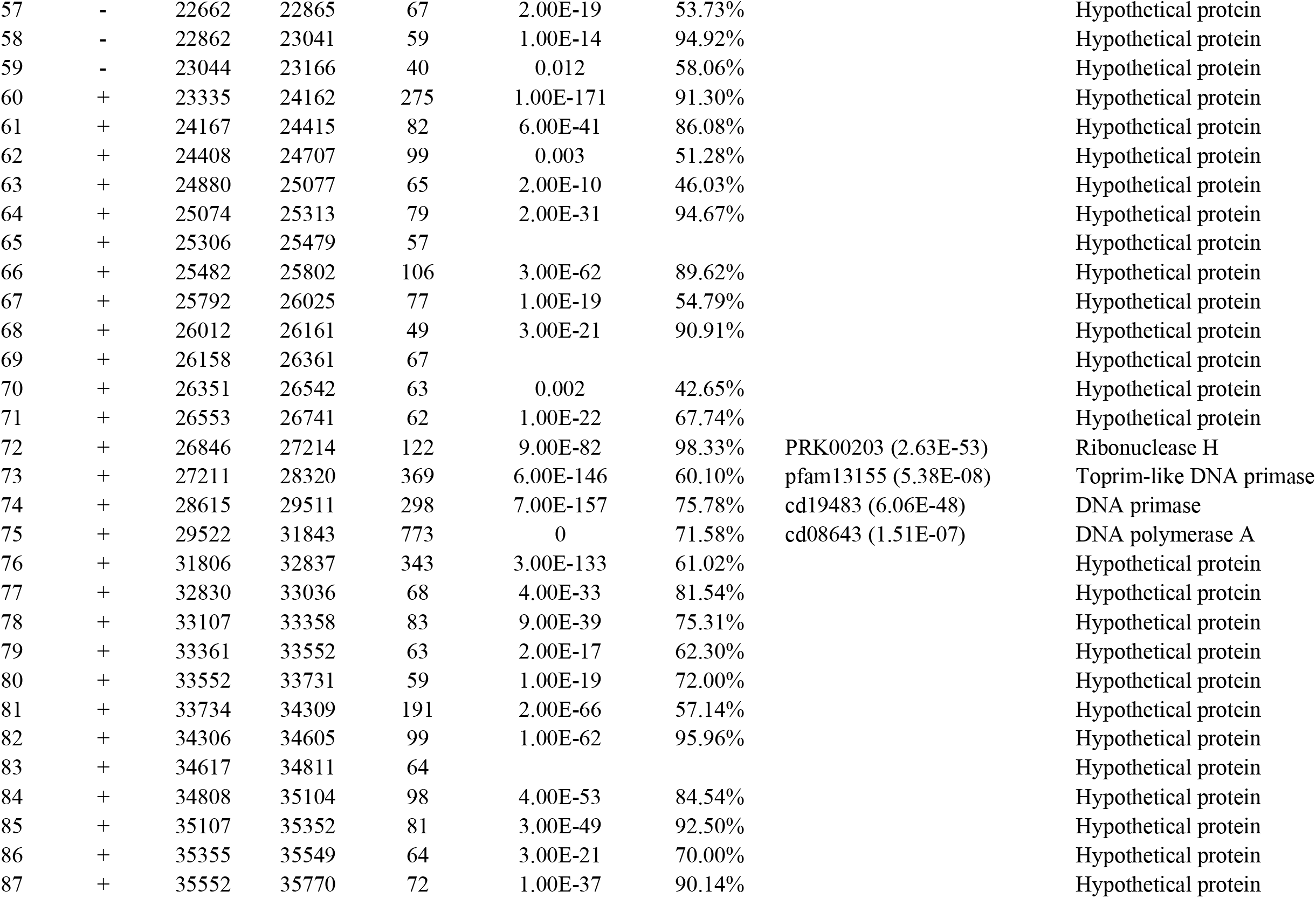
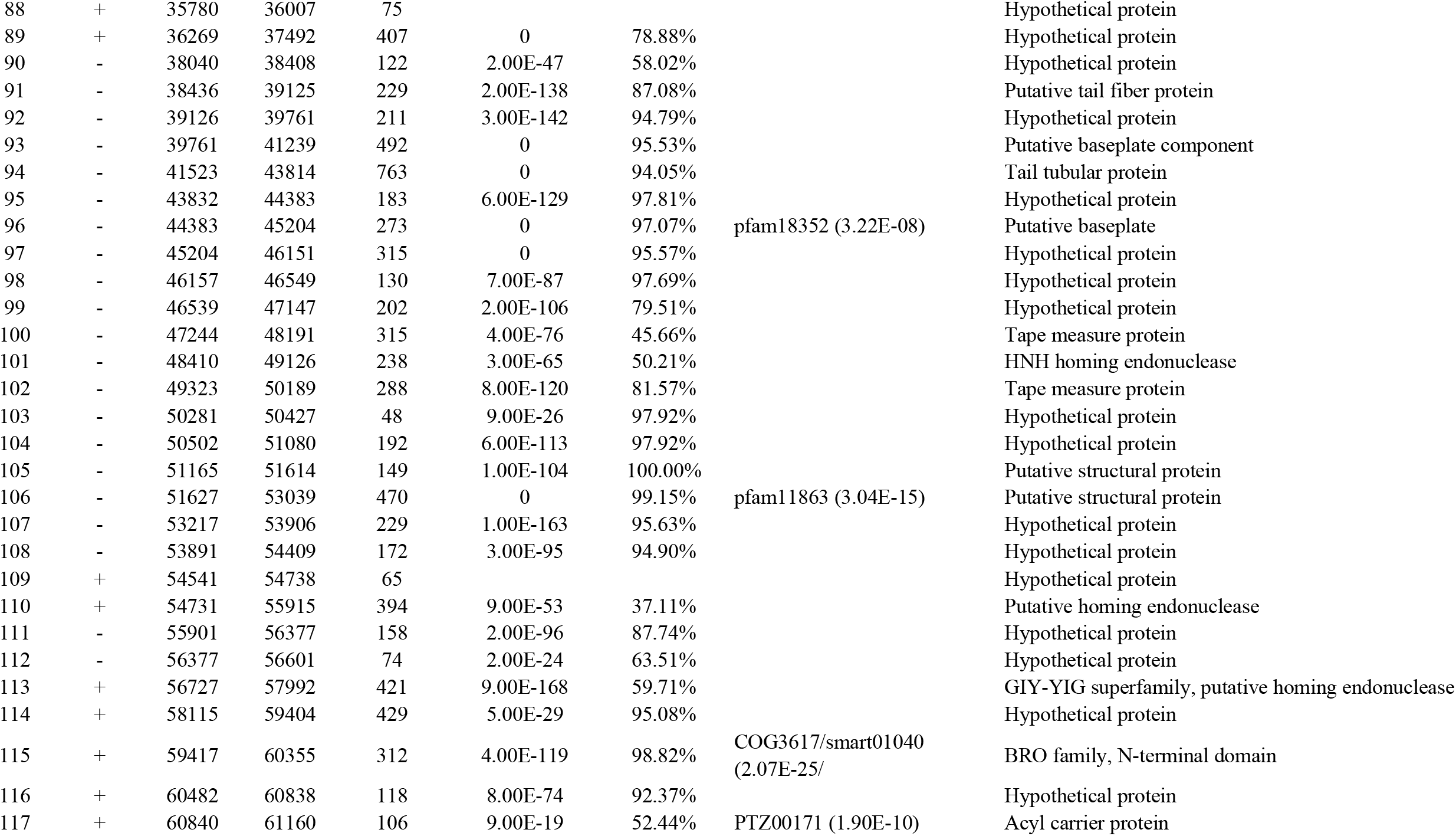
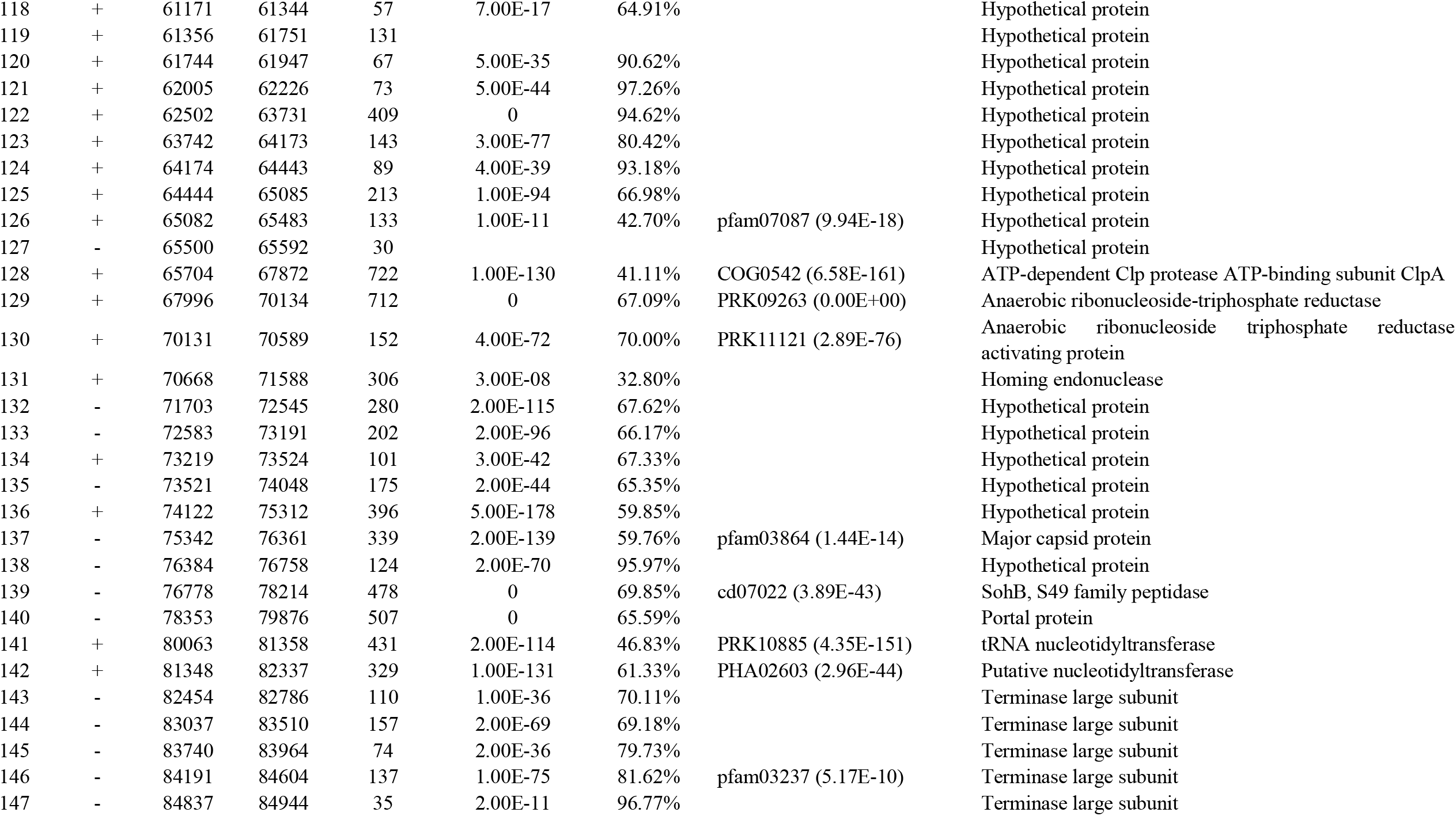
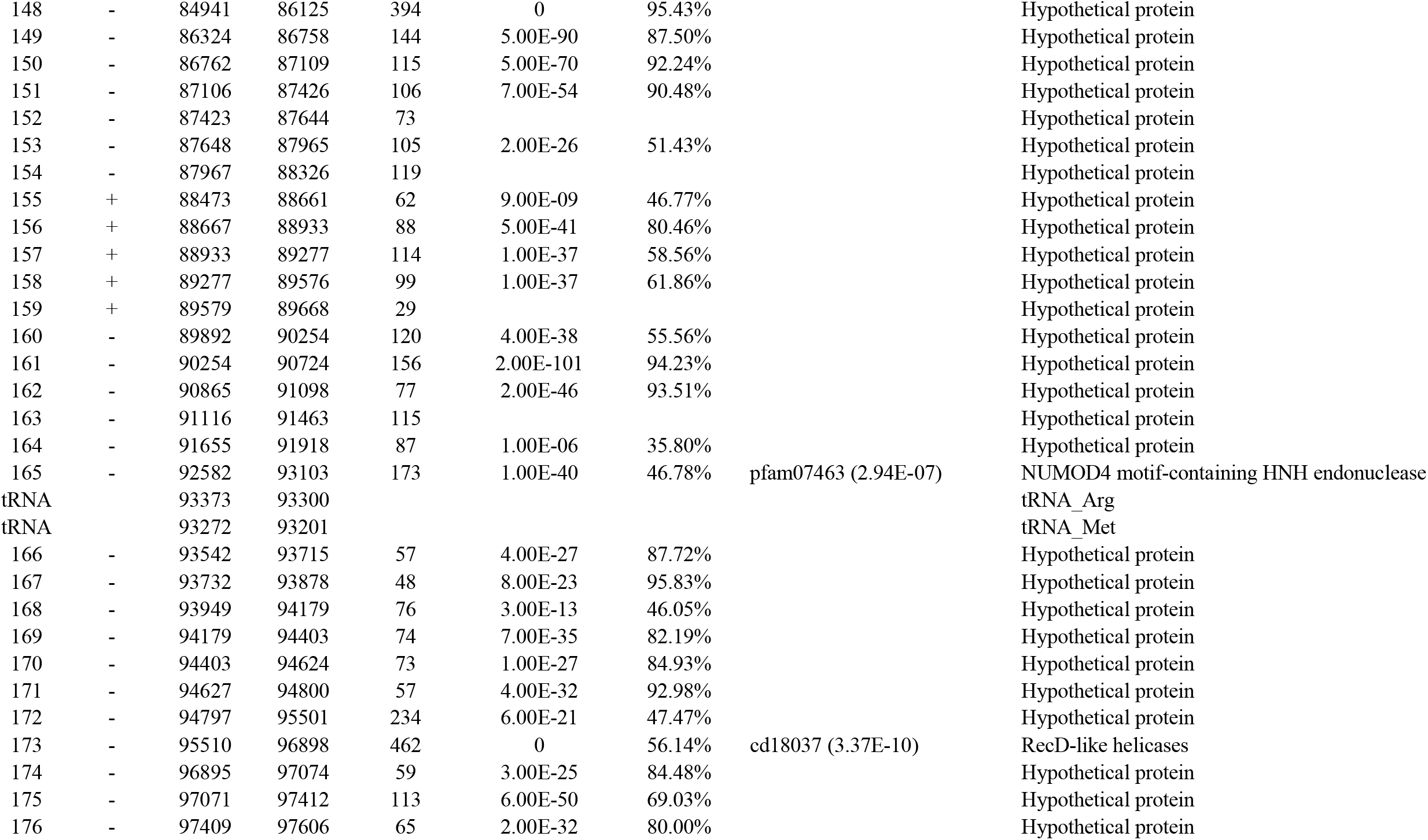
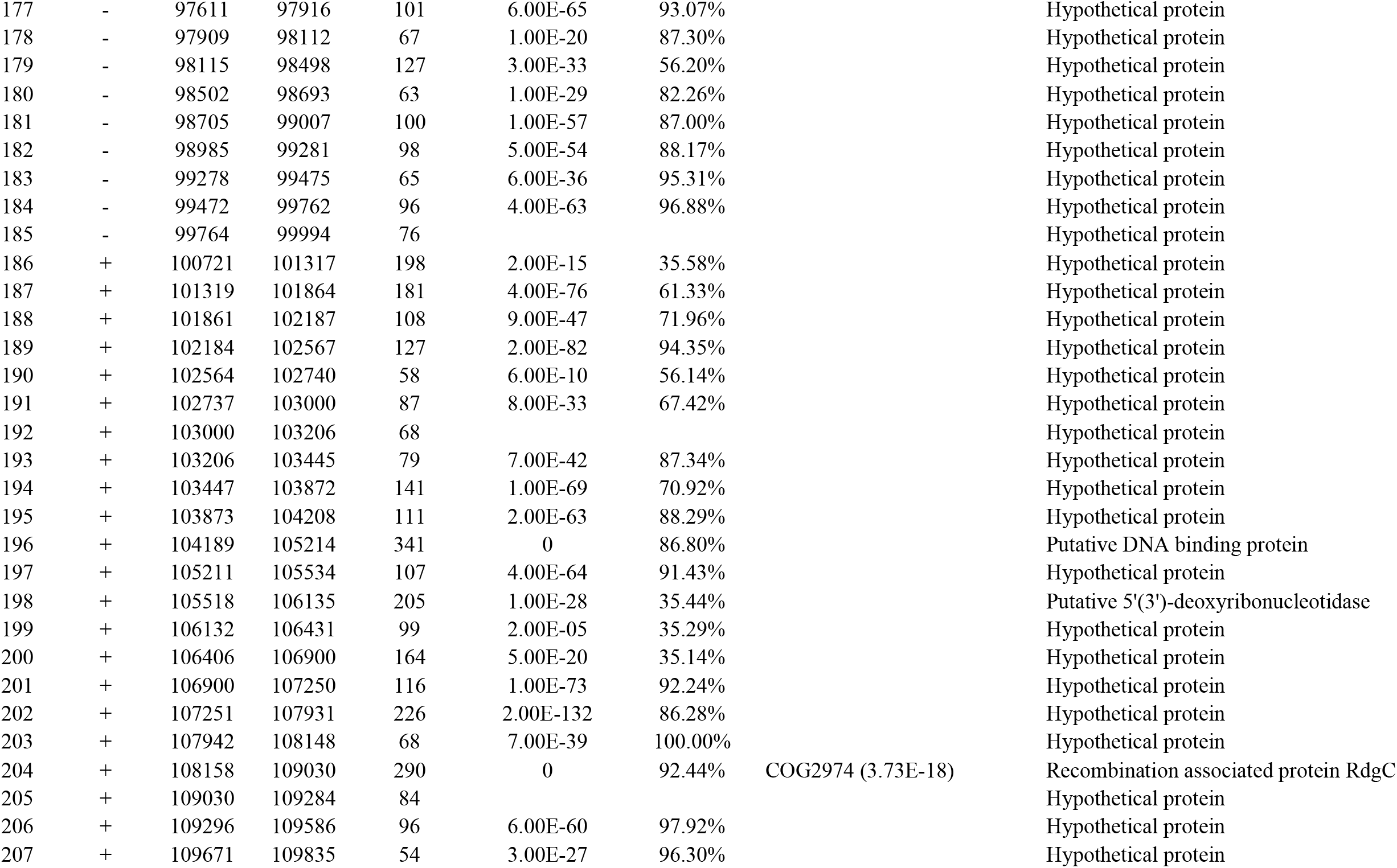
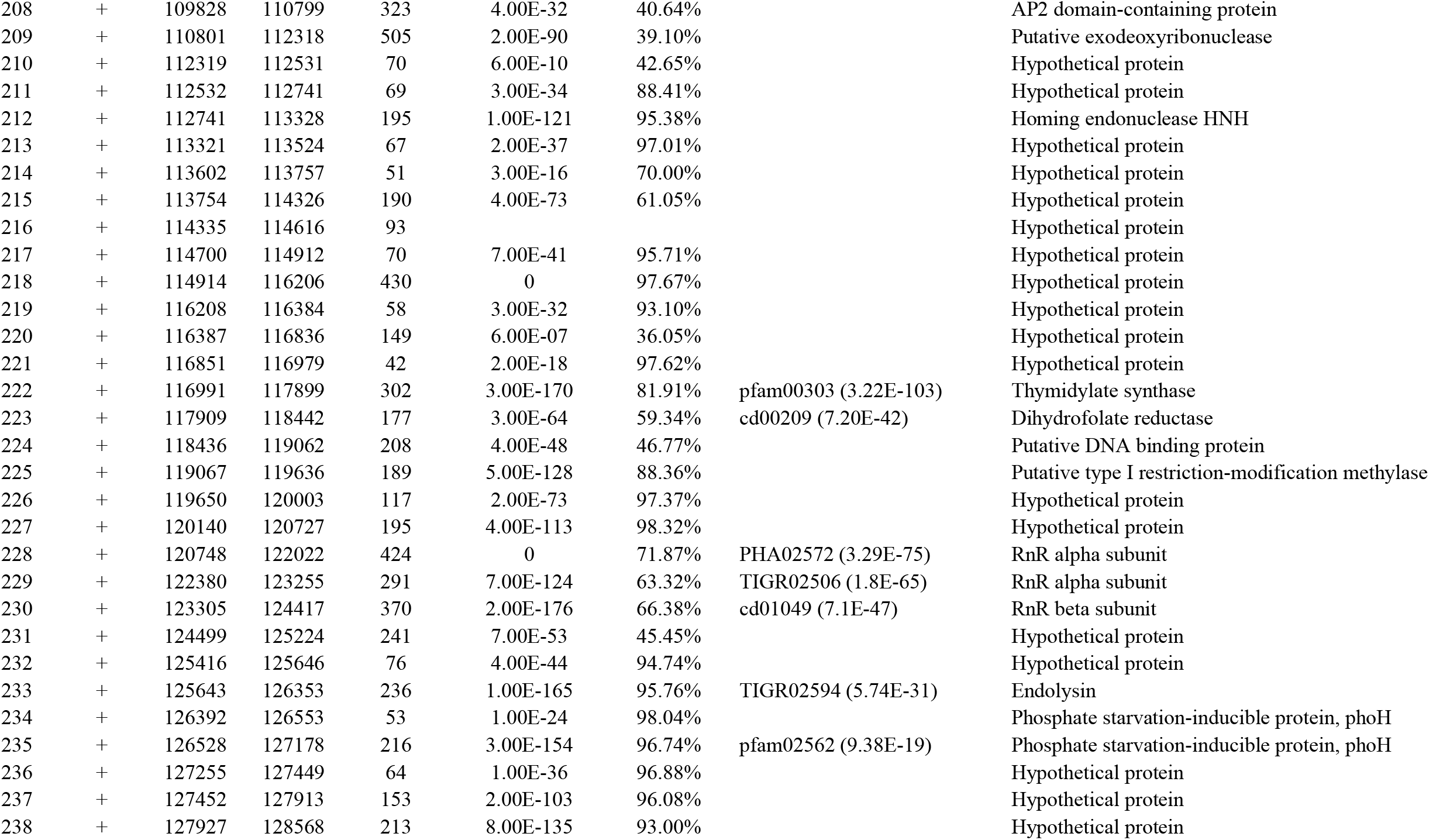
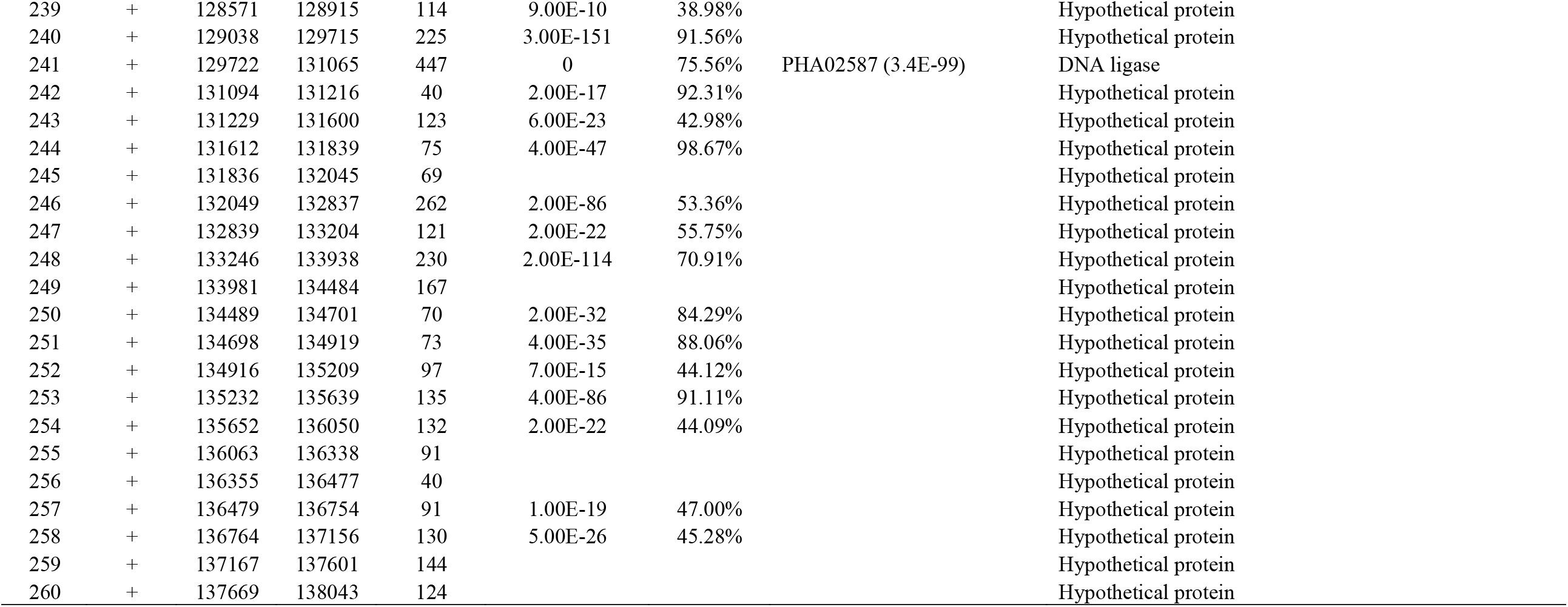
Predicted ORFs in the R16F genome.

**Table S3.** Related phages identified by vCONTACT2 (score > 1)

| **Phage** | **GenBank** | **Score** | **Morphology** |
| --- | --- | --- | --- |
| *Vibrio* phage helene 12B3 | NC_021067 | 57.7337 | Myovirus |
| *Vibrio* phage eugene 12A10 | NC_029067 | 49.1551 | Myovirus |
| *Aeromonas* phage 2-L372D | NC_048769 | 13.2045 | Myovirus |
| *Aeromonas* phage 4_L372D | NC_048771 | 13.5277 | Myovirus |
| *Aeromonas* phage 2_L372X | NC_048770 | 13.4260 | Myovirus |
| *Aeromonas* phage 4_L372XY | NC_048772 | 13.0604 | Myovirus |
| *Enterobacteria* phage phi92 | NC_023693 | 5.1463 | Myovirus |
| *Klebsiella* phage ZCKP1 | NC_047994 | 3.5774 | Myovirus |
| *Escherichia* phage ESCO13 | NC_047770 | 3.3215 | Myovirus |
| *Escherichia* phage vB_EcoM_Schickermooser | NC_048196 | 3.2685 | Myovirus |
| *Escherichia* phage ESCO5 | NC_047776 | 4.8704 | Myovirus |
| *Escherichia* phage phAPEC8 | NC_020079 | 2.3294 | Myovirus |
| *Pectobacterium* phage phiTE | NC_020201 | 1.5967 | Myovirus |
| *Escherichia* phage slur16 | NC_028248 | 1.0244 | Myovirus |
| *Escherichia* phage V18 | NC_041937 | 1.0063 | Myovirus |
| *Escherichia* phage Av-05 | NC_025830 | 4.8183 | Myovirus |
| *Erwinia* phage Hena1 | NC_048828 | 2.8524 | Myovirus |
| *Erwinia* phage pEp_SNUABM_01 | NC_048807 | 1.4785 | Myovirus |
| *Raoultella* phage Ro1 | NC_048682 | 6.6279 | Myovirus |
| *Klebsiella* virus KpS8 | NC_048873 | 3.6915 | Myovirus |
| *Klebsiella* phage vB_KpnM_KB57 | NC_028659 | 2.7575 | Myovirus |
| *Proteus* phage Mydo | NC_048741 | 3.6341 | Myovirus |
| *Klebsiella* phage vB_KpnM_BIS47 | NC_048656 | 3.6723 | Myovirus |
| *Salmonella* phage PVPSE1 | NC_016071 | 5.3238 | Myovirus |
| *Salmonella* phage SSE121 | NC_027351 | 5.3692 | Myovirus |
| *Escherichia* phage 4MG | NC_022968 | 3.5028 | Myovirus |
| *Cronobacter* phage vB_CsaM_GAP31 | NC_019400 | 3.5400 | Myovirus |
| *Rheinheimera* phage vB_RspM_Barba19A | NC_048190 | 1.0529 | Myovirus |
| *Rheinheimera* phage vB_RspM_Barba8S | NC_048188 | 1.1008 | Myovirus |
| *Vibrio* phage ICP1 | NC_015157 | 54.4929 | Myovirus |
| *Vibrio* phage PWH3a-P1 | NC_020863 | 185.9040 | Myovirus |
| *Vibrio* phage qdvp001 | NC_029057 | 238.3224 | Myovirus |
| *Vibrio* phage 11895-B1 | NC_020843 | 55.8402 | Myovirus |
| *Shewanella* sp. phage 1/4 | NC_025436 | 9.6671 | Myovirus |
| *Shewanella* sp. phage 1/40 | NC_025470 | 20.2678 | Myovirus |
| *Pseudoalteromonas* phage H101 | NC_029094 | 12.8149 | Myovirus |
| *Pseudoalteromonas* phage J2-1 | MF370964 | 32.0631 | Myovirus |

